# Slap restricts oncogenic Src-family kinase signaling to maintain colonic epithelial homeostasis

**DOI:** 10.64898/2026.01.05.697659

**Authors:** Dana Naim, Zouheir Houhou, Florent Cauchois, Kevin Espie, Valérie Simon, Yvan Boublik, Francina Langa Vives, Zeinab Homayed, Conception Paul, Morgan Maillard, Michael Hahne, Julie Pannequin, Julie Nguyen, Audrey Sirvent, Serge Roche

## Abstract

Src-family kinases (SFKs) regulate proliferation in colonic epithelial cells (CECs), but the mechanisms that restrain their activity remain poorly defined. We identify Src-like adaptor protein (SLAP), a negative regulator of receptor tyrosine kinase signaling, as a key suppressor of SFK activity in the colon. Constitutive and inducible epithelial-specific Slap deletion using a villin-CreERT2 model increases CEC proliferation and accelerates tumorigenesis in the azoxymethane/dextran sodium sulfate (AOM/DSS) model. Slap deficiency also enhances SFK-dependent expansion of normal and tumor-derived colonic organoids. Mechanistically, we identify the receptor tyrosine kinase EPHB2 as a critical upstream activator of SFKs and a direct target of SLAP-mediated regulation. Loss of Slap increased EphB2 protein abundance and tyrosine phosphorylation and enhanced its association with active SRC. Pharmacological inhibition of EPHB2 suppressed SRC activation and reversed the hyperproliferative phenotype induced by Slap deficiency. Together, these findings uncover a non-genetic mechanism driving SFK activation during colonic transformation and establish SLAP as a tumor suppressor that constrains oncogenic EPHB2–SFK signaling in the colonic epithelium.

## INTRODUCTION

The membrane-anchored tyrosine kinase (TK) SRC plays important roles in intestinal homeostasis and tumorigenesis. In particular, SRC and other members of its family (i.e., FYN and YES) drive intestinal stem/progenitor cell proliferation, tissue regeneration and tumorigenesis in Drosophila and mouse models (Cordero et al., 2014; Imada et al., 2016; Kohlmaier et al., 2015; Sirvent et al., 2020). SRC is also an oncogene in human colorectal cancer (CRC). Aberrant SRC expression and activity in CRC (50% of CRC patients) is a marker of poor clinical prognosis, a promoter of therapeutic resistance and a potent driver of metastasis (Sirvent et al., 2012; Summy and Gallick, 2003; Yang et al., 2022). This tumour activity is associated with cancer stem cell and epithelial-to-mesenchymal transition features. However, SRC is rarely mutated in CRC and the mechanisms underlying its oncogenic function remain elusive (Irby et al., 1999; Sirvent et al., 2012; Summy and Gallick, 2003). Consistently, SRC inhibitors developed for the clinic have failed in CRC due to ineffective signalling inhibition and inadequate patient selection (Daud et al., 2012; Parseghian et al., 2017; Reddy et al., 2015). Elucidating the mechanisms underlying this persistent signalling may reveal essential non-genetic mechanisms by which TKs promote tumorigenesis and allow the development of effective TK-based therapies in CRC.

During evolution, TK activities have come under the tight control of small adaptor proteins that exert negative regulatory functions (Naudin et al., 2016). The best example is the SOCS regulation of JAK/STAT inflammatory pathways. We discovered that SRC signalling is under the control of such an inhibitory mechanism through the membrane-anchored adaptor SLAP (Src-Like Adaptor Protein), which display high sequence homology with SRC N-terminus (Manes et al., 2000; Pandey et al., 1995; Roche et al., 1998; Sirvent et al., 2008). Mechanistically, SLAP promotes degradation or prevents SRC binding to upstream receptors, resulting in inhibition of SRC signaling (Dragone et al., 2006a, 2006b; Kazi and Rönnstrand, 2012; Myers et al., 2006, 2005; Roche et al., 1998; Sirvent et al., 2008; Wybenga-Groot and McGlade, 2015). Slap-deficient mice revealed an essential function for this adaptor in controlling lymphocyte development and activation, where it is highly expressed (Sosinowski et al., 2001). We additionally reported an unsuspected tumour suppressor function of SLAP in CRC (Naudin et al., 2014). Notably, SLAP was found to be abundantly expressed in the intestinal epithelium. SLAP levels were reduced in 50% of CRC samples analysed, and SLAP silencing in experimental CRC models promoted tumour formation and liver metastasis. Mechanistically, SLAP promotes degradation of SRC signalling RTK EPHA2 (Naudin et al., 2014) and destabilization of the mTORC2 complex, resulting in AKT signaling inhibition (Mevizou et al, 2025). Although this observation suggests an important mechanism in the control of SRC function in the colon, genetic evidence for this mechanism is lacking. Here, we developed Slap-deficient mice models showing that Slap regulates SFK colonic epithelial function during homeostasis and tumorigenesis.

## MATERIALS AND METHODS

### Generation of genetically modified mouse models

All mouse experiments were conducted in strict accordance with the European Community guidelines (86/609/EEC) and the French National Committee for the care and use of laboratory animals (87/848), complied with the ARRIVE guidelines, and were approved by the French Ministry of Higher Education, Research and Innovation (APAFIS#2022031511382008). All procedures were performed in the institutional animal facility (agreement #F3417216). Mice were housed in temperature-controlled ventilated cages (20–22 °C) under a 12 h light/dark cycle, with relative humidity maintained between 45% and 55%, and were kept under pathogen-free conditions. *Slap^tm1a(KOMP)^* mice were generated from JM8A1.N3 embryonic stem cells (C57BL/6N genetic background) produced by the trans-NIH KnockOut Mouse Project (KOMP) and obtained from the KOMP Repository (www.komp.org). Genotyping was performed by PCR analysis of genomic DNA isolated from mouse tail biopsies. The Slap ^tm1a(KOMP)^, Tg(CAG-flpo)1Afst, and Tg(Vil1-cre/ERT2)23Syr mouse lines, as well as compound lines, were maintained on a C57BL/6 background and bred in a specific opportunistic pathogen-free (SOPF) facility, and experiments were conducted in a specific pathogen-free (SPF) facility.

*Slap*-inactivated mice (*Slap -/-*) were generated by insertion of a *LacZ* reporter cassette into the Slap locus, resulting in functional gene inactivation. The *LacZ* cassette was flanked by FRT sites, allowing recombination upon crossing with FlpO deleter mice. FlpO-mediated recombination generated the recombined *Slap* allele, whereas FlpO-negative littermates retaining the unrecombined *LacZ* allele were used as controls.

Inducible *Slap* knockout mice were generated by crossing Slap floxed mice with CreERT2 transgenic mice. Tamoxifen treatment induced Cre-mediated excision of the floxed Slap allele, resulting in gene inactivation. To activate CreERT2, control and experimental mice received two intraperitoneal injections of tamoxifen (2 mg per injection).

### Mice genotyping

Mice tail biopsies underwent alkaline lysis and incubation at 92 °C for 20 minutes, followed by reaction neutralization. PCR amplification was conducted with the ready mix MyTAQRed mix (meridian). Resulting products were run on a 2% ethidium-bromide agarose gel (Sigma–Aldrich), using a 1 kb DNA ladder (Thermo Scientific), and imaged with a ChemiDoc MP system (Bio-Rad). The following primers were used: CSD-lacF: GCTACCATTACCAGTTGGTCTGGTGTC; CSD-neoF: GGGATCTCATGCTGGAGTTCTTCG; CSD-loxF: GAGATGGCGCAACGCAATTAATG; CSD-Sla-R: TCTCTGAGATGCCCCATTTTACCCC; CSD-Sla-ttR: GTTTCTTGGCACTGACTAGAGCAGG; CSD-Sla-F: GTTAACAACACACCTACCCCTTGGC.

### Chemical induction of tumorigenesis by AOM/DSS

Inducible epithelial SLAP knockout mouse model was treated with tamoxifen to induce Slap depletion. DSS and AOM/DSS treatment was performed essentially as described in (Nguyen et al., 2025; Paul et al., 2022). Briefly, 10 days following tamoxifen treatment, a first AOM injection was performed. Shortly after the first round of DSS at 2% was given to the mice for 7 consecutive days. A week later another AOM injection was performed followed by 2 other rounds of DSS treatments two weeks apart. The mice were then sacrificed at day 80. Colons or tumors were then excised for tumoroids formation, cryopreservation or fixation followed with IHC analysis.

### β-galactosidase activity

To visualize the activity of the Slap promoter driving lacZ expression, we used 5-bromo-4-chloro-3-indolyl-beta-D-galactopyranoside (Xgal, Sigma-Aldrich) as described in (Nguyen et al., 2025). Organs were cryopreserved and 10 μm tissue sections were prepared. Sections were treated with 0.5% glutaraldehyde for 10 min at room temperature followed by overnight incubation at 37°C in a staining solution containing 1 mg/ml X-gal, 5 mM potassium ferricyanide (K3Fe(CN)6), 5 mM potassium ferrocyanide (K4Fe(CN)6) and 2 mM MgCl2, 0.1% Triton in PBS-1X. The synthetic substrate X-gal gives an insoluble blue precipitate when cleaved by β-galactosidase. Samples were washed in PBS-1X for 5 min and rinsed briefly with dH2O before mounting with aqueous mounting medium (Sigma-Aldrich).

### Immunohistochemistry of paraffin embedded tissues

Fluorescent and bright-field immunohistochemistry on paraffin-embedded tissue Intestinal tissues were fixed in neutral-buffered formalin for 24 hours at RT, embedded in paraffin and tissue sections (4 μm) were prepared with a microtome (MICROM). Sections were incubated in successive baths of xylene and ethanol and antigen retrieval was performed by boiling the sections with 10 mM sodium citrate (pH 6.4) or Tris-EDTA buffer (pH 9) for 20 minutes. Tissues were treated with TBS-1 × 2% serum 0.1% Triton to block non-specific binding. Samples were then incubated with primary antibodies overnight at 4 °C. In the case of Bright-field immunohistochemistry, sections were treated with BIOXALL (Vector laboratories) for 10 minutes before blocking and incubation with primary antibodies. Secondary reagent staining was revealed using DAB (Vector Laboratories), and sections were counterstained with hematoxylin (Sigma-Aldrich) and then mounted with aqueous mounting medium (Sigma-Aldrich). As for fluorescence immunohistochemistry, sections were incubated in the presence of secondary antibodies conjugated to Alexa-488 or cyanin3 (Jackson ImmunoResearch Laboratories) and Hoechst (2 µg/mL, Sigma–Aldrich) and mounted with aqueous mounting medium (Sigma-Aldrich). For histological analysis, the tissue sections were deparaffinized and stained with hematoxylin, eosin, and Alcian blue.

### RNA in situ hybridization

ISH for Lgr5 was performed with the RNAscope FFPE assay kit (Bio-techne) according to the manufacturer’s instructions. Briefly, 4 mm formalin-fixed, paraffin embedded tissue sections were pretreated with heat and protease digestion and then hybridized with a target probe for Lgr5 (Probe - Hs-LGR5-C2, Biotechne). Thereafter, a fluorophore signal amplification system was hybridized to the target probe.

### RNA extraction and qPCR

Total RNA was extracted using RNeasy Plus Mini Kit columns (Qiagen). First-strand cDNA synthesis was performed with 1 µg of purified RNA using SuperScript VILO cDNA synthesis KIT (Thermo Fisher Scientific) according to the manufacturer’s instructions. qRT-PCR experiments were performed using LightCycler 480 SYBR Green I Master (Roche Diagnostics) on the LightCycler 480 system using the following primers EHPB2_F 5’-CGGCTGCATGTCCCTCATC-3’, EHPB2_R 5’-GTCCCCGTTACAGTAGAGCTT-3’; GAPDH_F 5’-TCTCCTCTGACTTCAACAGCGAC-3’ & GAPDH_R 5’-CCCTGTTGCTGTAGCCAAATTC-3. Relative expression was determined using the threshold cycle relative quantification method. Data were normalized to the expression levels of GAPDH.

### ALDH analysis

The ALDH assay was performed as described in (Nguyen et al., 2025) and according to the protocol of the ALDEFLUORTM kit (#01700, Stemcell). FACS analysis of was preformed through the Novocyte ACEA 2 fluorescence-activated cell sorting flow cytometer and data were analyzed using NovoExpress software.

### Antibodies and reagents

Antibodies used in for biochemistry: anti-Myc (#2276S, CST), anti-EphB2 D2X2I (#83029, CST), anti-EphB2 (for IHC, # AF467 R&D Systems), anti-FLAG (M2 antibody, Sigma-Aldrich), anti-HA (Invitrogen, # 26183), anti-Actin (Sigma-Aldrich, #A2228), anti-pTyr 4G10 (gift from P. Mangeat, CRBM, Montpellier, France), anti-SLAP clone C-19 (Santa Cruz Biotechnology, sc-1215). SFK activity was assessed using an anti-phospho-SRC antibody (Invitrogen, PA5-97366 for WB, and #2101 from CST for IHC), which also cross-reacts with other p-SFKs, revealed increased SFK activity. All antibodies utilized in IHC were used according to supplier protocols. SFK activities were revealed with the help of SignalStain® Boost IHC Detection Reagent. Inhibitors used in this study: SRCi eCF506 (MedChemExpress, #HY-112096), EPHB2 inhibitor ALW-II-49-7 (EPHB2i, MedChemExpress, #HY-18833), Pan Eph inhibitor ALW-II-41-27 (pan-EPHi, MedCehmExpress, # HY-18007). SLAP-FLAG (WT and SH3*SH2* mutant, mut) constructs were described in (Naudin et al., 2014). EPHB2-myc construct was from (Narayanan et al., 2023). EPHB2 Y594F/Y602F mutant (YF mutant) was obtained by QuickChange Site-Directed Mutagenesis Kit (Agilent) using the following oligonucleotides: 5’-gaccccaggcatgaagatctTcatcgatcctttcacctTcgaggaccccaacgaggcag-3’ and 5’-ctgcctcgttggggtcctcgAaggtgaaaggatcgatgAagatcttcatgcctggggtc-3’. Other reagents: Advanced DMEM/F12 (Gibco), B27 supplement (ThermoFischer Scientific), N2 supplement (ThermoFischer Scientific), GlutaMAX (ThermoFischer Scientific), Hepes (ThermoFischer Scientific), N-acetyl cysteine (Sigma-Aldrich), CHIR-99021 (Tebu-BIO), Recombinant human R-spondin-1 (Peprotech). Tamoxifen (Merck group), Azoxymlethane (Sigma-Aldrich).

### Cell culture and transfections

HT29, SW620 and HEK293T cell lines were obtained from ATCC (Rockville, MD). HT29 and SW620 cells stably expressing SLAP (or mock as a control) were described in (Mevizou et al, 2025; Naudin et al., 2014). Cells were cultured at 37°C and 5% CO2 in a humidified incubator in Dulbecco’s Modified Eagle’s Medium (DMEM) GlutaMAX (Invitrogen) supplemented with 10% fetal calf serum (FCS), 100 U/ml of penicillin and 100 µg/ml of streptomycin, and were routinely tested (once a week) for Mycoplasma using MycoAlert Mycoplasma detection kit (Lonza, #LT07-318). Transient plasmid transfections in HEK293T cells were performed with the jetPEI reagent (Polyplus-transfection) according to the manufacturer’s instructions.

### Biochemistry

Co-immunoprecipitation and Western Blotting (WB) were described in (Mevizou et al, 2025; Naudin et al., 2014). Cells or isolated colonic crypts were lysed on ice using lysis buffer (20 mM Hepes pH 7.5, 150 mM NaCl, 0.5% Triton X-100, 6 mM β-octylglucoside), supplemented with complete mini EDTA-free protease and phosphatase inhibitor cocktail tablets (Roche). Protein lysates (15-35 µg) were resolved by SDS-PAGE, transferred to LF/PVDF membranes using the Trans-Blot® Turbo™ system (Bio-Rad), and incubated overnight at 4 °C with the appropriate primary antibody (1:1000). Membranes were then incubated for 45 minutes with the HRP-conjugated secondary antibody (Cell Signaling, 1:4000). Signals were detected using ECL Plus (Amersham Biosciences) and imaged with the Amersham Imager 600 (GE Healthcare Life Sciences). Signal quantification and analysis were carried out using ImageJ software. For SLAP-EPHB2 association experiments, HEK293T cells were co-transfected for 48 h with the indicated FLAG-tagged and EPHB2 constructs either WT or the YF EPHB2 mutant. Endogenous EPHB2 association experiments were performed in HT29 cells stably expressing SLAP-FLAG. Cell-lysates were subjected to SLAP-FLAG immunoprecipitation using anti-FLAG magnetic beads (#A3697, Pierce). EPHB2-myc was immunoprecipitated using MYC-Tag (9B11) mouse monoclonal antibody (#2276, CST) and protein G sepharose beads (Thermo Fisher Scientific).

### Organoids derived from colonic crypts

Inducible epithelial Slap knockout models were sacrificed 10 days after tamoxifen induction. Organoids were performed as described in (Nguyen et al., 2025). Briefly, the colons were collected and washed with PBS containing antibiotics. They were then cut open and further dissected into small pieces then washed with a wash buffer containing PBS 1x, penicillin streptomycin (Gibco), and Fungin (Invivo Gen) until supernatant is clear. These pieces were then incubated in a crypt isolation buffer on ice for 30 minutes (Wash buffer, 1% BSA fraction and 25 mM EDTA). Crypt fractions were then isolated after thorough pipetting, centrifuged at 300g, 4°C for 5 minutes. 500 crypts were resuspended in M1 medium mixture (Advanced DMEM/F12 supplemented with 1% Penstrep, 1% Fungin, 10 mM Hepes, 1% GlutaMAX, N-2, B-27, 1 Mm N-acetyl cysteine and Matrigel (Corning) (ratio 1:2) The mixture was plated in a 24 well plate. After Matrigel polymerization, 500 µL of M2 medium (M1 media supplemented with EGF (50 ng/mL, Bio-techne), Noggin (100 ng/mL, Stem cell technologies), R-spondin1 (500 ng/mL, Stem cell technologies), Y27 (10 µM, Sigma-Aldrich), CHIR-99021 (3 µM, Tebu-Bio) and WNT3a (50 n/ mL, Thermo Fisher Scientific) was added to each well containing or not the SRC inhibitor eCF506 (100 nM, Medchem express) and in other cases the EphB inhibitors EPHB2i (ALW-II-49-7, 200 nM) and the pan-EPH inhibitor pan-EPHi (ALW-II-41-27, 100 nM, Medchem express). For organoids derived from AOM/DSS transformed CECs, tumors were excised from mice treated with AOM/DSS at day 80. They were dissociated using the Tumor Dissociation Kit mouse (Miltenyi Biotec). The dissociation was the stopped by DMEM medium containing 10% SVF. The mixture was then centrifuged the pellet was resuspended in DMEM F12 without factors. Cells were counted and plated at almost 10,000 cells per well in a 24 well plate in a mixture containing 3D Matrigel. After 30 minutes of incubation a complete F12 medium was added to the solidified domes containing or not the SRC inhibitor eCF506 (100 nM) (Fraser *et al*, 2016).

### Tumoroids-derived from CRC cells

2 000 - 5 000 human CRC cells were resuspended in Advanced DMEM/F12 supplemented with 1% Penstrep, 2mM L-glutamine, N-2 and Matrigel (Corning) (1:2 ratio) prior to plating in 24-well plates. After polymerization of Matrigel, 0.5 mL Advanced DMEM/F12 supplemented with EGF (20 ng/ml, Biotechne) and FGF (10 ng/ml, Biotechne) was added. Tumoroids were cultured at 37°C and 5% CO2 in a humidified incubator.

### Microscopy and imaging

Histological slides were digitized using a Nanozoomer scanner (Hamamatsu) with a 40X objective and viewed with NDP.view2 software (Hamamatsu). Fluorescence images were captured using an AxioImager Z2 microscope (Zeiss) operated with Zen software. Image processing was performed in ImageJ.

### TCGA COAD analysis

Gene expression and clinical data from the TCGA colon adenocarcinoma (COAD) cohort were analyzed using GEPIA2 (http://gepia2.cancer-pku.cn/). Kaplan–Meier disease-free survival analyses were performed using the GEPIA2 Survival Analysis module, with patients stratified into high-and low-expression groups based on the median expression value. Stem cell-like signature scores were calculated using the GEPIA2 Signature Score module (Merlos-Suárez et al., 2011) from the indicated gene set, and associations with disease-free survival were assessed by log-rank testing.

### Statistical analysis

All analysis was performed using GraphPad Prism (9.3.1). Data are presented as the mean ± SEM. the Mann-Whitney test was used. The statistical significance level is illustrated with p values: *p≤0.05, **p≤0.01, ***p≤0.005, **** p≤0.0051(t-test).

## RESULTS

### Constitutive *Slap* inactivation increases CEC proliferation

To investigate SLAP function in the colon, we generated constitutive Slap-deficient mice by inserting a *LacZ* cassette into the Slap gene (*Slap-/-* mice) (Fig. 1A). β-galactosidase activity staining revealed a gradient of Slap expression toward the top of the crypts throughout the colonic epithelium (Fig. S1A). This expression pattern was confirmed at the protein level by immunofluorescence staining using a Slap-specific antibody (Fig. 1A). Loss of Slap led to increased colon thickness in 3-month-old mice, characterized by elongated crypts (Fig. 1A). This epithelial hyperplasia was accompanied by elevated colonic epithelial cell (CEC) proliferation (Fig. 1A) and an increased number of goblet cells (Fig. S1B), indicating that Slap regulates both CEC proliferation and differentiation.

**Figure 1.**
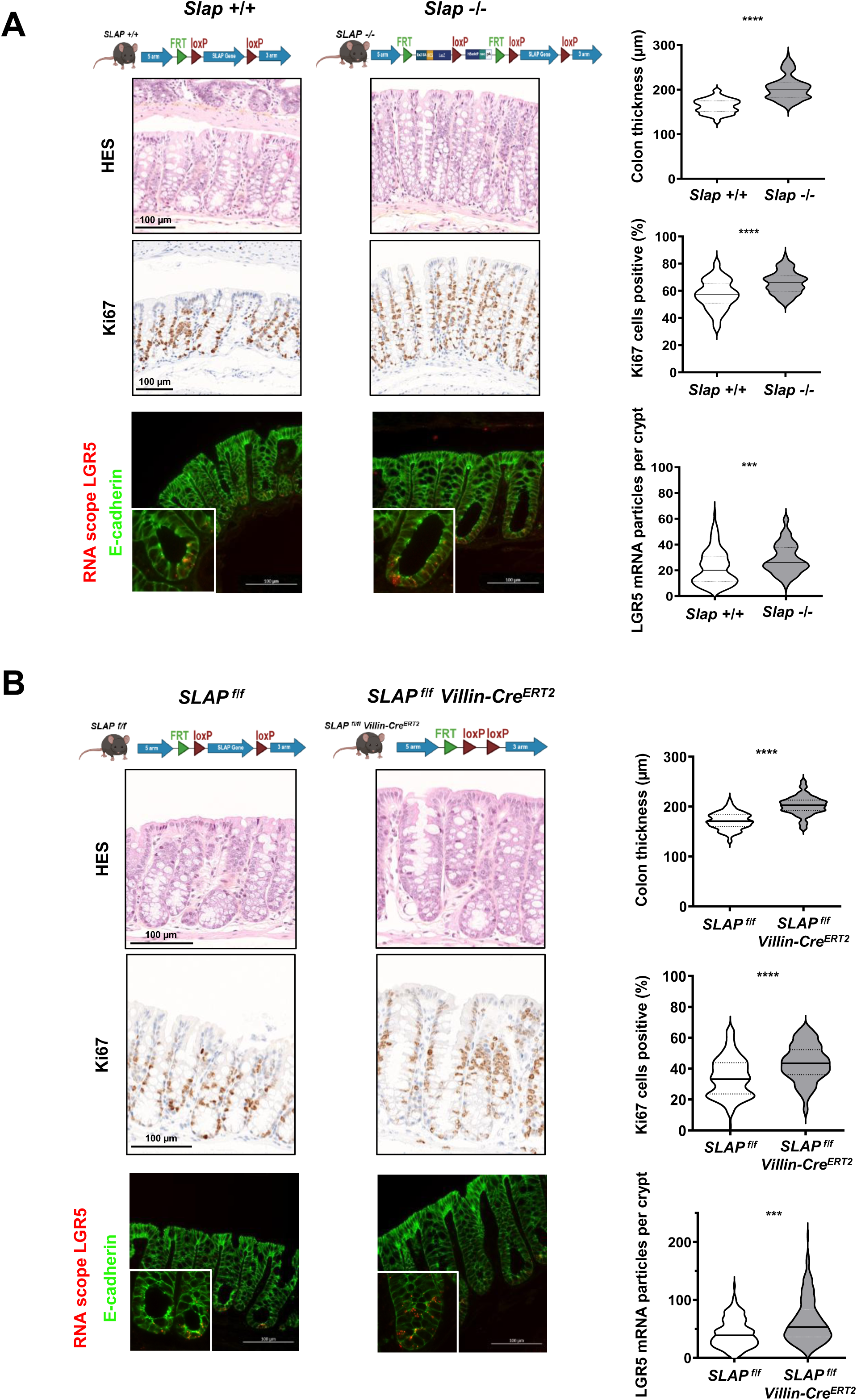
Murine Slap inactivation induces colonic epithelial hyperplasia. **(A)** Constitutive Slap deletion. Top: Hematoxylin and eosin (H&E) staining of transverse colon sections. Crypt thickness was measured and quantified as mean ± SEM from 25-40 crypts per mouse (n = 6 female mice per group). Middle: Immunohistochemistry (IHC) for the proliferation marker Ki67, with quantification of Ki67-positive colonic epithelial cells (CECs). (mean ± SEM from 20-40 crypts per mouse; n = 5-6 mice per group). Bottom: RNAscope analysis of *Lgr5* to identify colonic stem cells, combined with E-cadherin immunofluorescence to delineate crypt architecture. Inset, higher (4X) magnification highlighting *Lgr5* expression. *Lgr5* RNA particles were quantified and are shown as mean ± SEM from 20-40 crypts per mouse (n = 5-6 mice per group). ***p<0.001, ****p<0.0001 Mann–Whitney test. **(B)** Inducible epithelial-specific Slap deletion (villin-CreERT2). Top: H&E staining of transverse colon sections with quantification of crypt thickness (mean ± SEM from 25-40 crypts per mouse, n = 6 mice/group). Middle: IHC for Ki67 with quantification of Ki67-positive CECs (mean ± SEM from 20-40 crypts per mouse; n = 5-6 mice per group). Bottom: RNAscope analysis of Lgr5 combined with E-cadherin immunofluorescence; Inset, higher (4X) magnification highlighting *Lgr5* expression. Lgr5 RNA particles were quantified (mean ± SEM from 20-40 crypts per mouse, n = 5-7 mice per group). Mice were analyzed at 3 months of age, 10 days after tamoxifen induction. ***p<0.001, ****p<0.0001 Mann–Whitney test.

RNAscope analysis of Lgr5, a marker of colonic epithelial stem cells (CSCs) (Barker et al., 2007), showed increased expression at crypt bases in Slap-deficient mice (Fig. 1A), suggesting enhanced CSC traits. Given the role of SFKs in regulating colonic stem/progenitor proliferation (Cordero et al., 2014; Imada et al., 2016; Kohlmaier et al., 2015; Sirvent et al., 2020), we assessed whether Slap modulates SFK activity. Immunohistochemical analysis of an antibody detecting the activated form of Src (pSRC), which also cross-reacts with other p-SFKs, revealed increased SFK activity throughout the crypts of Slap-deficient mice (Fig. S1B). This suggests that SLAP plays a role in regulating intestinal epithelial homeostasis, in addition to its well-established function in lymphocyte development (Sosinowski et al., 2001).

### Cell-autonomous role of Slap in CEC proliferation

To determine if Slap functions in a cell-autonomous manner, we generated an inducible, intestinal-specific Slap-deficient mouse model by crossing *Slap flox/flox* mice with *villin-CreERT2* mice (*Slap ^f/f^ Villin Cre^ERT2^*). Tamoxifen treatment for 10 days efficiently deleted Slap in the epithelium, as confirmed by a marked reduction in Slap immunofluorescence staining in the intestinal epithelium, while resident immune cells retained Slap expression (Fig. S1B). Colonic Slap depletion recapitulated the phenotype of constitutive Slap deficient mice, including increased colon thickness, enhanced CEC proliferation (Fig. 1B), elevated goblet cell numbers (Fig. S1C), and increased CSC activity (Fig. 1B). To further investigate the cell-autonomous role of Slap in regulating CEC proliferation, we performed ex vivo analyses using isolated colonic crypts. Organoids derived from Slap-deficient epithelium were more numerous and larger than those from controls (Fig. 2A).

**Figure 2.**
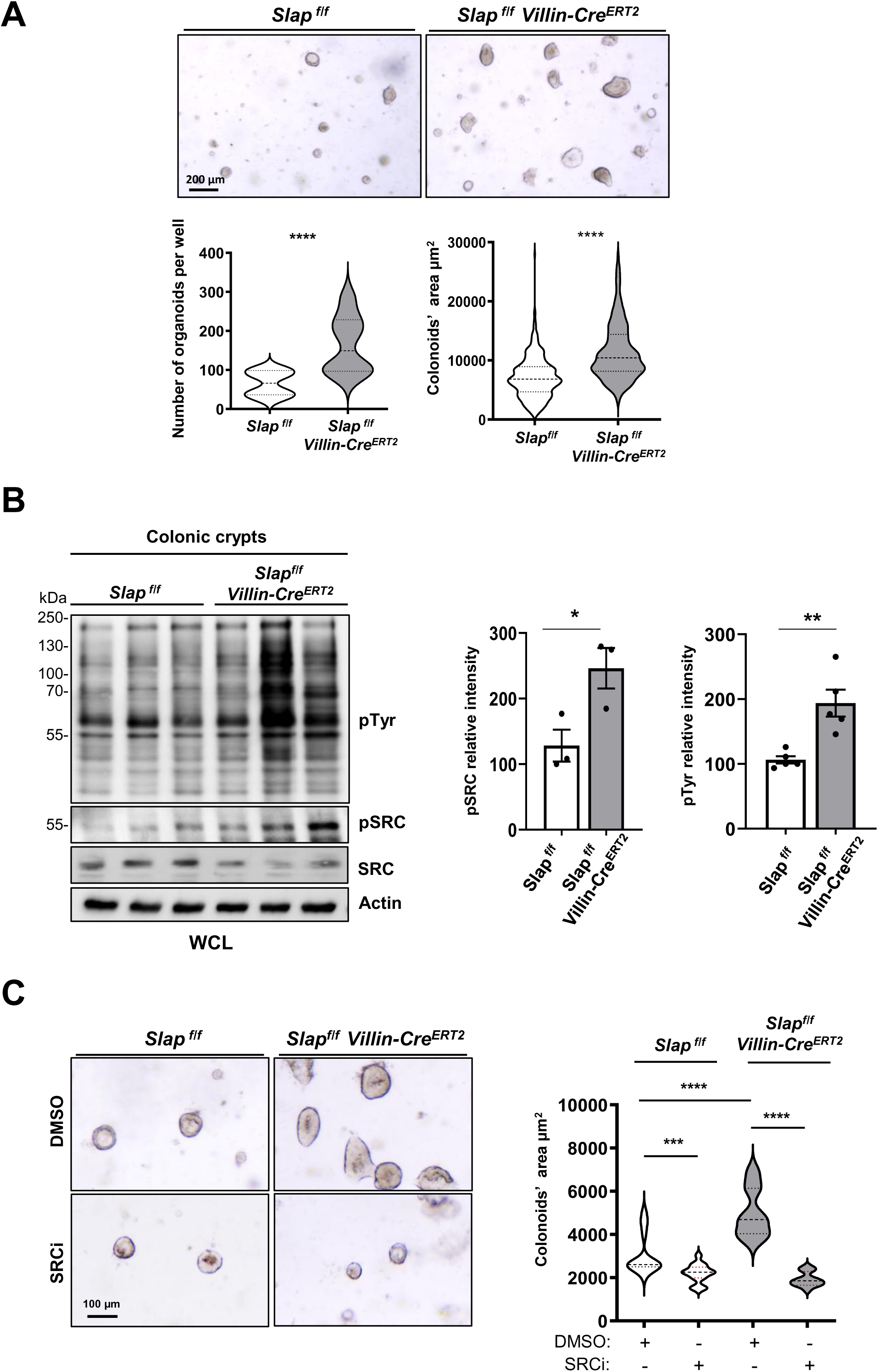
Intestinal Slap inactivation enhances colonic organoid development through SFK activation. **(A)** Slap-dependent colonic organoid development. Representative images of colon-derived organoids from *Slap ^f/f^* and *Slap ^f/f^ Villin-Cre^ERT2^* mice cultured in Matrigel for 2 days. Quantification of organoid number and area is shown as mean ± SEM from 100-200 organoids per mouse (n = 8 mice per group). ****p<0.0001 Mann–Whitney test. **(B)** Intestinal Slap deletion increases SFK activity and global protein tyrosine phosphorylation in colonic crypts. Representative immunoblots (right) and quantification (left) of phospho-SFK (pSRC) and total phospho-tyrosine levels in lysates from isolated colonic crypts of the indicated genotypes. Data are presented as mean ± SEM from n = 4-5 mice per group. *p<0.05; **p<0.001 t test. **(C)** Slap-dependent colonic organoid expansion requires SFK activity. Representative images and quantification of colon-derived organoids from *Slap ^f/f^* and *Slap ^f/f^ Villin-Cre^ERT2^* mice cultured for 2 days in the presence or absence of the SRC-family kinase inhibitor eCF506 (100 nM). Data are shown as mean ± SEM from 100-200 organoids per mouse, n = 4 mice per group. ***p<0.001, ****p<0.0001 Mann–Whitney test.

Consistent with a role for Slap in restraining SFK signaling, Slap-deficient colonic crypts exhibited a 2-3-fold increase in SFK activity and global cellular protein tyrosine phosphorylation (pTyr) levels (Fig. 2B). An increase in protein tyrosine phosphorylation was also observed in organoids derived from Slap-deficient CECs and was reduced following treatment with the selective SFK inhibitor eCF506 (SRCi) (Fraser et al., 2016). Consistently, SRCi treatment decreased pSRC levels, confirming effective SFK inhibition (Fig. S2). Functionally, the enhanced growth of Slap-deficient organoids was abolished by eCF506 treatment, demonstrating that the proliferative phenotype was SFK-dependent (Fig. 2C). Together, these findings establish SLAP as a cell-autonomous suppressor of SFK-driven CEC proliferation.

### Inducible Slap inactivation in the colon promotes colon tumorigenesis

We next evaluated Slap’s role in colon tumorigenesis using the azoxymethane/dextran sulfate sodium (AOM/DSS) model of colitis-associated cancer (CAC), in which a single injection of the carcinogen AOM is followed by repeated cycles of DSS-induced colonic inflammation, leading to the development of colorectal tumors that recapitulate key features of inflammation-driven CRC (De Robertis et al., 2011) (Fig. 3A). In the inducible Slap knockout model, Slap deletion was initiated by tamoxifen (TAM) administration 10 days before AOM treatment and maintained by a second TAM administration at day 45 to ensure sustained Slap inactivation throughout tumor development (Nguyen et al., 2025). Under these conditions, Slap-deficient mice developed significantly more tumors and exhibited increased SFK activity within tumors compared with control animals (Fig. 3B). Furthermore, tumoroids generated from these mice also showed enhanced growth, which was abolished by SFK kinase inhibition (Fig. 3C), further supporting an SFK-dependent oncogenic mechanism. Interestingly, pharmacological SFK inhibition had no effect on tumoroids derived from control AOM/DSS-treated mice, consistent with previous reports that AOM/DSS-induced carcinogenesis involves oncogenic RAS mutations (Takahashi & Wakabayashi, 2004) and chronic inflammation. These findings suggest that while RAS-driven transformation can occur independently of SFK activity, Slap loss unleashes a parallel, SFK-dependent oncogenic pathway. The human relevance of this SLAP anti-oncogenic function was confirmed by modulating SLAP expression in CRC cells. In human SLAP-low HT29 and SW620 cell-lines, SLAP overexpression inhibited aldehyde dehydrogenase (ALDH) activity (Fig. S3A), a well-established marker of CRC stem cells, as well as tumoroid growth (Fig. S3B). Together, these results establish SLAP as a critical suppressor of SFK-driven colon tumorigenesis and cancer stem cell activity.

**Figure 3.**
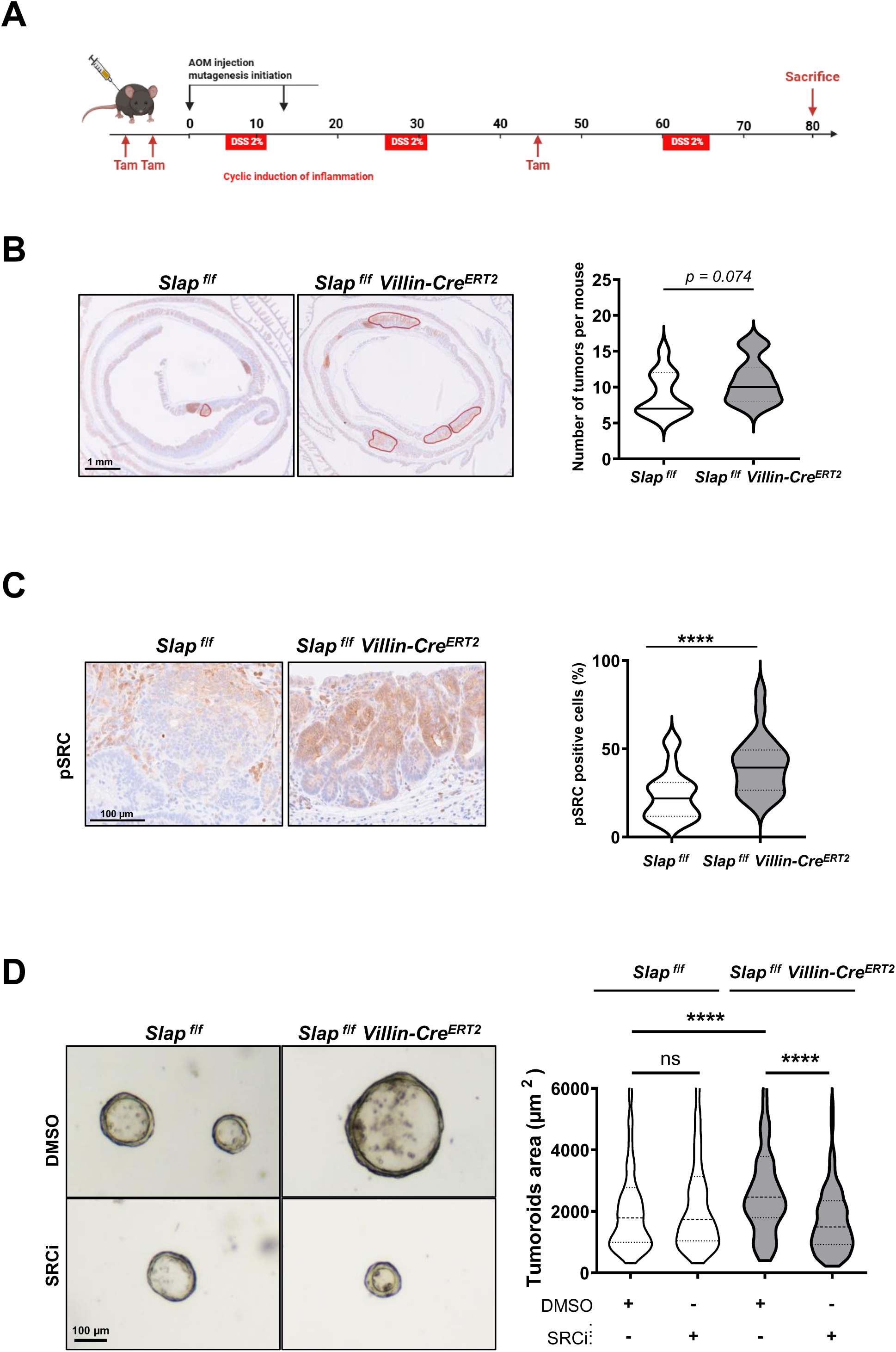
Inducible Slap intestinal inactivation drives colon tumorigenesis and promotes SFK-dependent tumoroid growth. **(A)** Experimental workflow of AOM/DSS-induced colon tumorigenesis. Mice received tamoxifen, followed 10 days later by a single azoxymethane (AOM) injection to induce mutagenesis and one cycle of dextran sodium sulfate (DSS) administered for 7 consecutive days. A second AOM injection was then performed, followed by two additional DSS cycles (7 days each). Mice were sacrificed at day 80. **(B)** Epithelial Slap loss accelerates colon tumor development. Left, representative H&E-stained sections showing transformed regions. Right, quantification of epithelial lesion area (mean ± SEM; n = 11-15 mice per group. Statistical analysis is shown (Mann–Whitney test). **(C)** Slap depletion increases SFK activation in colonic tumors. Left, representative immunohistochemistry for phosphorylated SFK (pSRC) in transformed epithelium. Right, quantification of p-SFK–positive cells per mm² (mean ± SEM; n = 7-9 mice per group; ****p < 0.0001, Mann–Whitney test). **(D)** Tumoroids derived from *Slap ^f/f^* and *Slap ^f/f^ Villin-Cre^ERT2^* mice were cultured in 3D Matrigel in the presence or absence of SRC inhibitor (SRCi) and imaged at day 3. Tumoroid area (µm²) was quantified following SRCi treatment (mean ± SEM; 150–250 organoids per mouse, n = 4 mice per group; ns: p>0.05; ****p < 0.0001, Mann–Whitney test).

### Slap target EphB2 to restrict SFK proliferative function in CEC

To investigate the Slap-dependent mechanism driving this phenotype, we hypothesized that Slap targets an upstream SFK activator within the RTK family that regulates colonic stem cells and CEC proliferation. Among RTKs expressed in colonic crypts, EphB2 and EphB3 are highly enriched in the CSC compartment and promote kinase-dependent CEC proliferation (Batlle et al., 2002; Genander et al., 2009; Holmberg et al., 2006). SFKs have also been reported as interactors and downstream effectors of EPHB2 signaling (Chang et al., 2024; Georgakopoulos et al., 2006; Leroy et al., 2009; Pasquale, 2024; Zisch et al., 1998). Supporting this model, Western Blotting analysis revealed elevated EphB2 protein levels in isolated Slap-deficient colonic crypts (Fig. 4A), which was further confirmed by IHC analysis showing increased EphB2 expression throughout the colonic crypts of Slap-deficient mice (Fig. 4B). This effect was recapitulated in CRC cells where enforced SLAP expression in SLAP-low HT29 cells decreased EPHB2 protein abundance. Interestingly, SLAP overexpression did not alter EPHB2 mRNA levels (Fig. S3A), in accordance with a posttranslational mechanism involved in this process, as reported for EPHA2 (Naudin et al., 2014). Co-expression experiments in HEK293T cells revealed that EPHB2 associates with SLAP (Fig. 4C), and this interaction required the SLAP SH2 and SH3 domains, as it was reduced with a SLAP mutant harboring inactive point mutations in the SH2 and SH3 domains (SLAP-mut) (Fig. 4C)(Mevizou et al., 2026). The interaction also depended on phosphorylation of the juxta-membrane Tyr596 and Tyr602, which mediates SRC-SH2 binding (Zisch et al., 1998), because mutation of these residues to phenylalanine (YF EPHB2) diminished SLAP-EPHB2 binding (Fig. 4C). Endogenous EPHB2 similarly interacted with SLAP in SLAP-expressing HT29 cells, confirming this association in CRC cells (Fig. S4B). Consistent with enhanced EphB2 signaling, isolated Slap-deficient colonic crypts displayed increased EphB2 tyrosine phosphorylation together with elevated levels of associated phosphorylated SFKs (Fig. 4A). Collectively, these findings identify EPHB2 as a novel SLAP interactor and target, supporting a role for SLAP in the regulation of EPHB2/SFK signaling.

**Figure 4.**
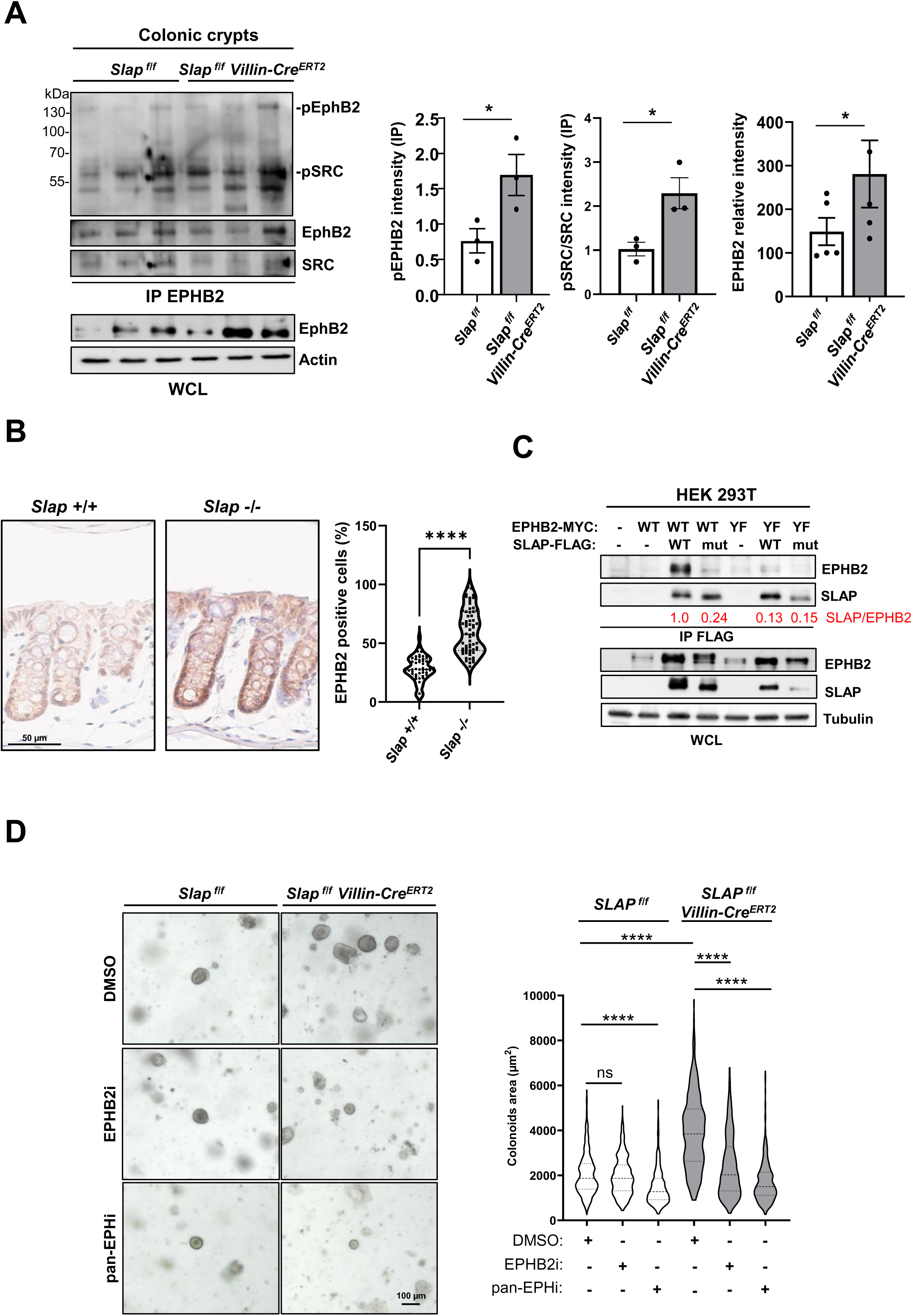
Slap targets EphB2 to restrict SFK proliferative signaling in CEC. **(A)** Intestinal Slap deletion increases EphB2 expression, EphB2 tyrosine phosphorylation, and its association with active SRC (pSRC) in colonic crypts. Left, quantification; right, representative immunoblots of EphB2 levels, EphB2 tyrosine phosphorylation, and associated pSRC following EphB2 immunoprecipitation from isolated colonic crypt lysates of the indicated mouse strains. Mean ± SEM; n = 3–5 mice per group; *p < 0.05, Student’s t-test. **(B)** IHC analysis of EphB2 expression in colonic crypts from control and Slap-deficient mice. Left, representative images; right, quantification (mean ± SEM; n = 3 mice per group; ****p < 0.0001, Mann–Whitney test). **(C)** SLAP–EPHB2 interaction in HEK293T cells. Co-immunoprecipitation of the indicated SLAP-FLAG constructs (wild-type, WT; or SH3*SH2* mutant, SLAP mut) and EPHB2-MYC constructs (wild-type, WT; or Y596F/Y602F mutant, YF) transfected into HEK293T cells. Expression levels of SLAP and EPHB2 and relative quantification of SLAP-EPHB2 interaction are shown (representative example of three independent experiments). **(C)** Slap-dependent colon organoid expansion requires EphB2 activity. Representative images and quantification of colon-derived organoids from *Slap ^f/f^* and *Slap ^f/f^ Villin-Cre^ERT2^* mice treated with the indicated EPH inhibitors (EPHB2i: 200 nM, pan-EPH2i: 100 nM, or DMSO control) and analyzed at day 2 (mean ± SEM; 150-200 organoids per mouse; n = 4 mice per group; ns: p>0.05; ****p < 0.0001, Mann–Whitney test).

We next asked whether EphB2 contributes to the Slap-dependent regulation of colonic organoid growth by inhibition of EphB2 kinase activity with the selective inhibitor ALW-II-49-7 (EPHB2i) (Choi et al., 2009; Noberini et al., 2012; Zhang et al., 2025). While EPHB2i had no effect on the growth of wild-type organoids, it markedly suppressed the enhanced growth observed in Slap-deficient organoids (Fig. 4C). Treatment with the pan-EPH kinase inhibitor ALW-II-41-27 (pan-EPHi) (Amato et al., 2014; Choi et al., 2009; Ferrao Blanco et al., 2024), produced a similar reduction (Fig. 4C), supporting a role for EphB2 kinase activity in the Slap-dependent phenotype. Of note, EPHB2i did not alter the kinase activity of SRC expressed in HEK293T cells in contrast to EPHB2, arguing against a direct off-target effect on SFK (Fig. S3B). Moreover, pEPHi reduced both the elevated phosphotyrosine levels and SFK activation induced by Slap deletion in colonic organoids (Fig. S2), demonstrating that SFK hyperactivation in Slap-deficient CECs is dependent upon EphB2 signaling. Together, these data show that Slap limits colonic organoid growth by restricting EphB2-dependent activation of Src and reveal an EphB2-SFK signaling axis as a key driver of the hyperproliferative phenotype associated with Slap-deficiency. (Fig. S4C). Notably, a similar mechanism was observed in human CRC cells. Pharmacological inhibition of EPHB2 reduced tumoroid growth in a CRC model expressing low levels of SLAP, whereas this inhibitory effect was largely abolished upon SLAP overexpression. Importantly, a SLAP mutant (SLAP mut) that is unable to interact with EPHB2 fails to suppress tumoroid growth and restores sensitivity to EPHB2 inhibition, indicating that the tumor-suppressive activity of SLAP depends on its interaction with EPHB2 (Fig. S5A-C). Collectively, these findings indicate that SLAP restrains the tumor-promoting activity of EPHB2 and suggest that the SLAP–EPHB2–SRC signaling axis is conserved between normal CECs and CRC.

Finally, we sought additional evidence supporting SLAP-EPHB2 signaling in CRC patients. We observed a weak but significant inverse correlation between SLAP expression and the colorectal CSC-like activity score in TCGA tumors (Fig. S5A). In addition, although expression of EPHB2, SLAP, or the SLAP-related gene SLA2 alone was not associated with disease-free relapse in microsatellite-stable (MSS) colorectal cancer patients (Fig S5B), co-expression of SLAP and SLAP2 with EPHB2 was associated with improved prognosis (Fig 5D). Notably, this association was specific to MSS tumors, which are characterized by low immune infiltration, and was not observed in microsatellite instability (MSI) CRC, which display high immune infiltration (Fig. 5D). Together, these findings suggest that SLAP expression in tumor cells may contribute to the regulation of EPHB2-dependent CSC signaling in CRC.

**Figure 5.**
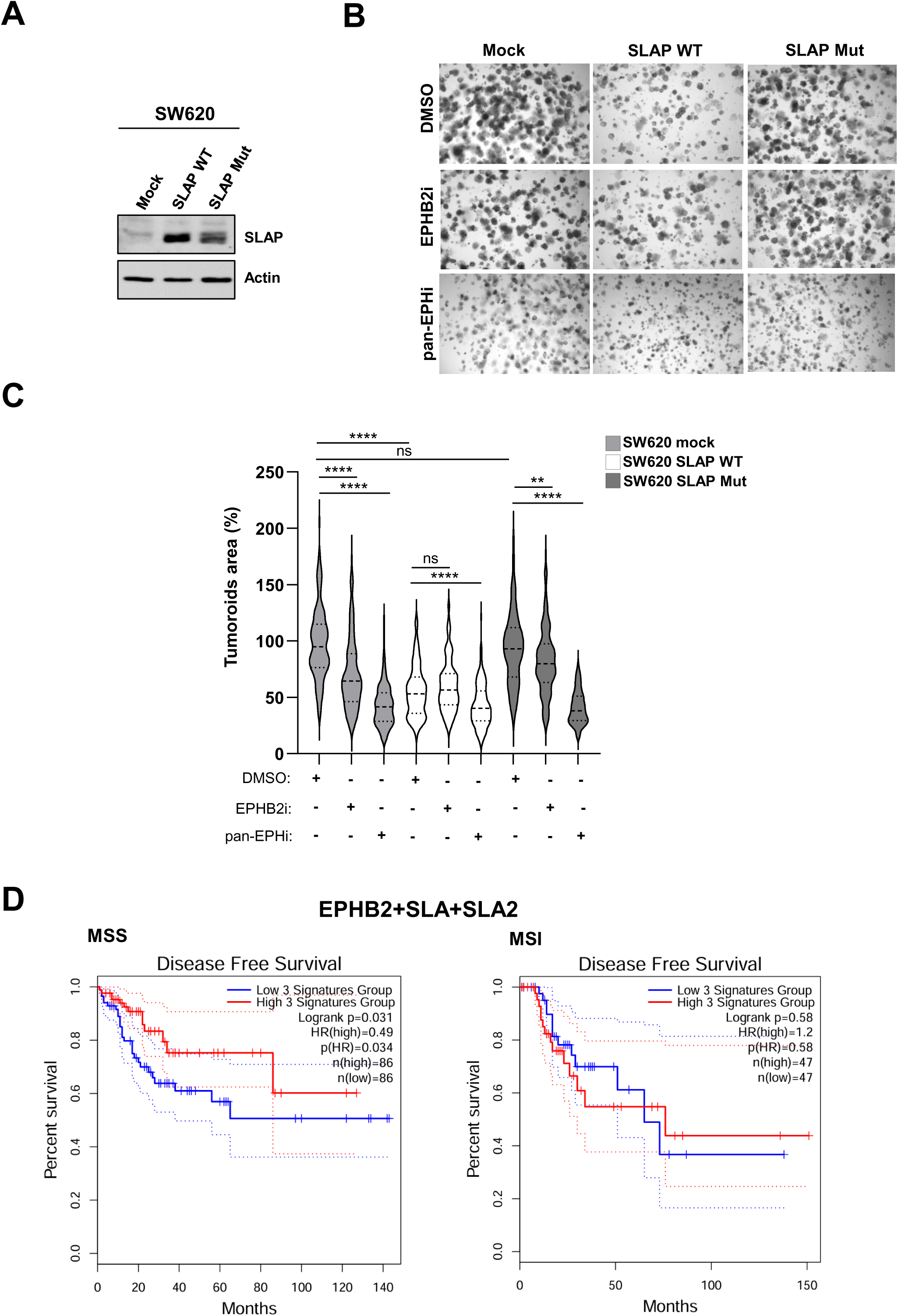
SLAP suppresses EPHB2-dependent tumoroid development of CRC cells and is associated with improved prognosis in MSS CRC. **(A)** SLAP expression in SW620 CRC cells transduced with empty vector (mock), wild-type SLAP, or an EPHB2-binding-defective SLAP mutant (SLAP Mut). **(B)** Representative tumoroid images from SW620 cells expressing mock, SLAP, or SLAP m and treated with indicated EPHB2 inhibitors (EPHB2i: 200 nM, pan-EPH2i: 100 nM, or DMSO control). **(C)** Quantification of tumoroid size from (B) (mean ± SEM; 150-200 tumoroids; n = 3; ns: p>0.05; **p<0.001; ***p<0.0005; ****p < 0.0001, Mann–Whitney test). **(D)** Disease-free relapse analysis of TCGA MSS and MSI CRC patients stratified according to combined EPHB2, SLAP, and SLA2 expression. High SLAP/SLA2 and EPHB2 co-expression is associated with improved prognosis in MSS patients.

## DISCUSSION

SFKs are potent drivers of intestinal stem cell proliferation and CRC, yet how their activity is restrained in normal epithelium remains unclear. Here, we identify the adaptor protein SLAP as a key epithelial-intrinsic regulator of SFK signaling in the colon. Epithelial loss of Slap increases proliferation, expands the stem cell compartment, and elevates SFK activation. Complementary organoid studies demonstrate that this requirement is cell autonomous. Importantly, pharmacologic SFK inhibition fully rescues the hyperproliferative phenotype of Slap-deficient organoids and tumoroids, revealing a crypt-intrinsic proliferative program that is SFK-dependent and normally constrained by Slap.

Mechanistically, our findings identify EPHB2 as a critical upstream activator of SRC signaling that is negatively regulated by SLAP. EPHB2 is highly expressed in intestinal stem and progenitor cells, where it contributes to crypt homeostasis and colorectal tumorigenesis (Batlle et al., 2005; Genander et al., 2009; Holmberg et al., 2006). We show that genetic loss of Slap results in elevated EPHB2 expression, increased SRC activation, and enhanced epithelial proliferation, whereas pharmacological inhibition of EPHB2 suppresses SRC activation and completely reverses the growth advantage of Slap-deficient organoids. Furthermore, EPHB2 inhibition selectively impairs the growth of CRC tumoroids expressing low levels of SLAP, supporting the existence of a conserved SLAP-EPHB2-SRC signaling axis in both normal and transformed intestinal epithelium.

Although the precise molecular mechanism remains to be established, SLAP may restrain EPHB2 signaling through several non-mutually exclusive mechanisms. SLAP has been shown to promote the ubiquitin-dependent downregulation of multiple RTKs through interactions with E3 ubiquitin ligases, including CBL family proteins, UBE4A, and UBE3C (Mevizou et al., 2025; Myers et al., 2006; Naudin et al., 2014; Sirvent et al., 2008; Zutshi et al., 2024). Alternatively, SLAP may limit productive SRC recruitment and activation downstream of EPHB2 through its SH2 and SH3 domains (Roche et al., 1998). Future studies will be required to determine how SLAP controls EPHB2 abundance and signaling output in intestinal stem cells. In addition, other RTKs known to activate SFKs, including members of the EGFR family (Cordero et al., 2014; Wong et al., 2012), may also contribute to the phenotype associated with SLAP loss. Moreover, Slap loss may affect EphB2’s role in cell positioning (Genander et al., 2009; Holmberg et al., 2006), contributing further to excessive proliferation.

Together, these findings establish SLAP as an epithelial tumor suppressor that limits SFK-driven stem cell proliferation and reveal a non-genetic mechanism of SRC oncogenic activation in CRC involving RTK hyperactivation. (Leroy et al., 2009). More broadly, they suggest that SLAP loss may define CRC subsets with heightened dependence on SFK signaling, offering opportunities for therapeutic stratification and refinement of SRC-targeted interventions.

## ACKNOWLEDGEMENTS

We thank the PCEA, ZEFI, Centre d’Ingénierie Génétique Murine and RHEM platforms animal facilities, immunohistochemistry analyses, Montpellier Rio Imaging (MRI) platform, Valérie PINET, Ethan ALBERT, Hahne’s and Pannequin’s teams, and our colleagues for discussion and technical support.

## AUTHORS CONTRIBUTION

All authors contributed extensively to the work presented in this paper. Experimental analysis and Data acquisition: DN, ZH, FC, VS, KE, YB, ZH, MM, JN, JP and AS. Genetical modified mouse models: AFL, AS, DN, CP and MH. Project supervision: SR and AS. Funding acquisition and Writing of the paper: SR.

## DATA AVAIBILITY

The data that support the findings of this study are available from the corresponding authors upon reasonable request.

---

**Figure S1.**
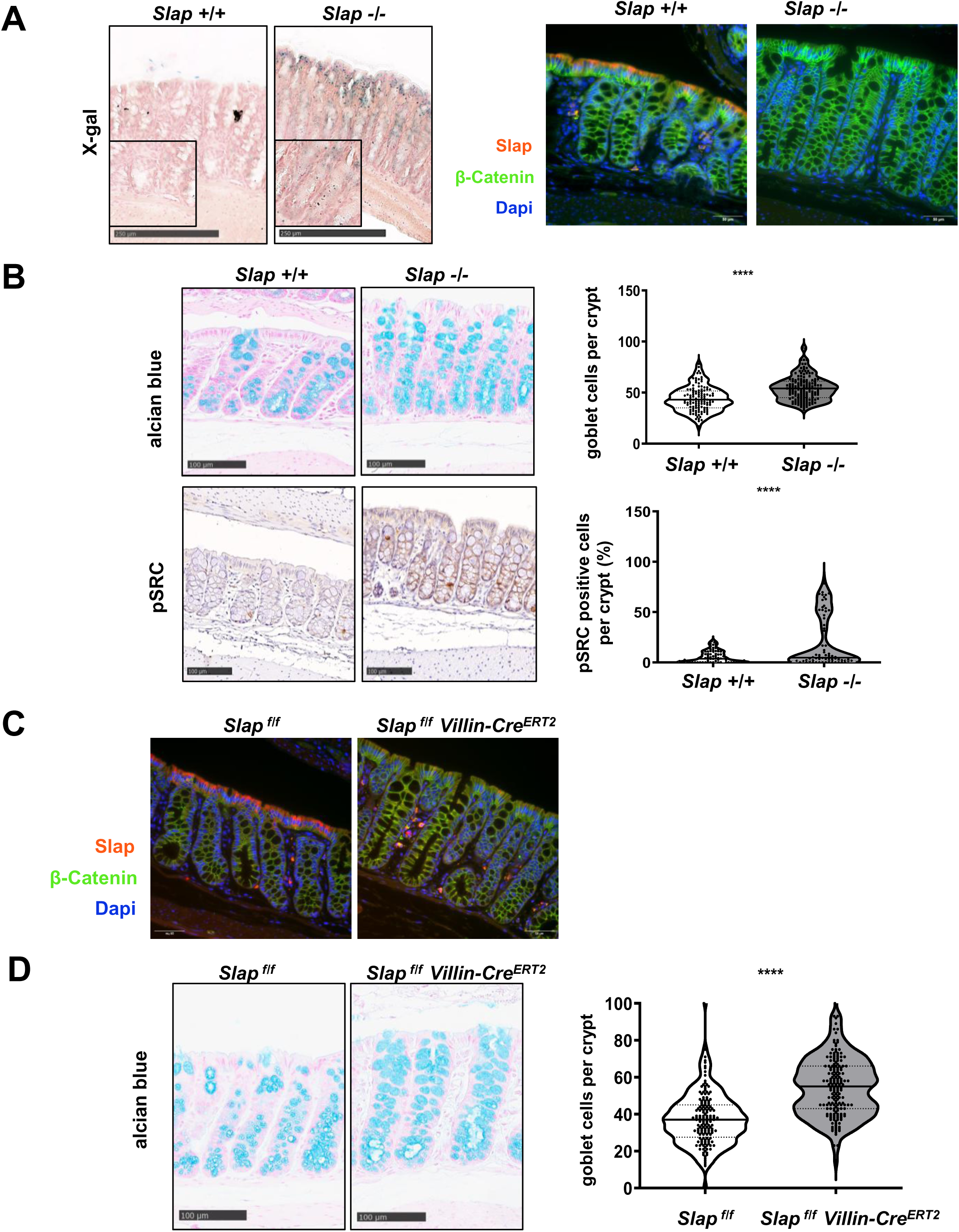
Slap inactivation increases goblet cell numbers and SFK activity in the colon. (**A**) Left: X-Gal staining of Slap expression. Insert: 4X magnification highlighting Slap expression at the base of colonic crypts. Right: IF staining for Slap (red), nuclei (blue), and β-catenin (green). (**B**) Slap inactivation increases goblet cell numbers and SFK activity. Top: Alcian blue staining of goblet cells with quantification. Bottom: IHC for p-SRC and its quantification.(**C**) Inducible Slap inactivation increases goblet cell numbers. Top: IF staining of Slap (red), nuclei (blue), and β-catenin (green) in the colon after 10 days of tamoxifen treatment. Bottom: Alcian blue staining of goblet cells and quantification.

**Figure S2.**
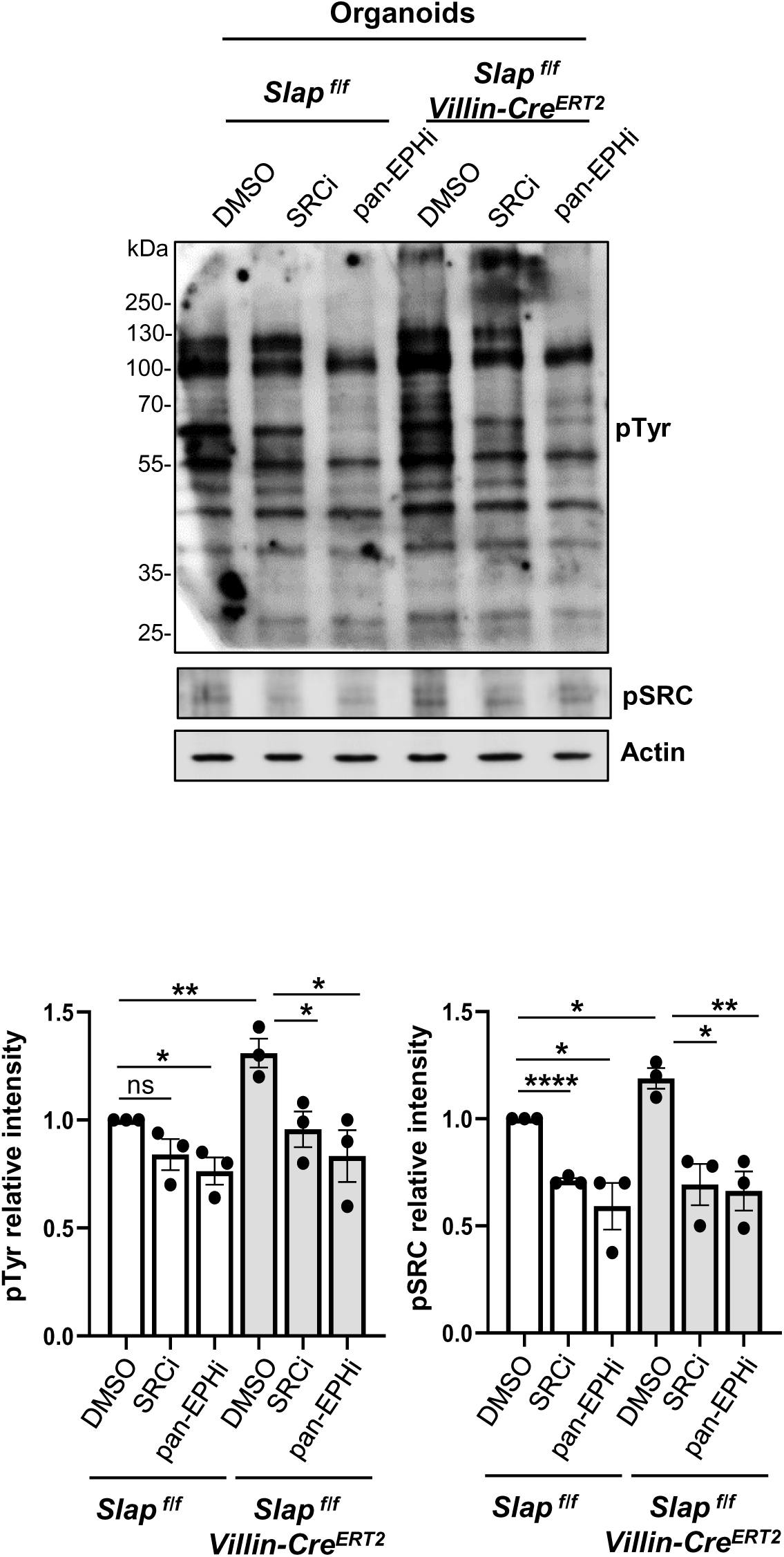
EPH-dependent phosphotyrosine accumulation and SFK activation in Slap-deficient colonic organoids. pTyr and activated SFKs (pSRC) levels in WT and Slap-deficient colonic organoids under the indicated conditions (EPHi and SRCi: 100nM for 3hrs); top: presentative example; bottom: quantification (mean ± SEM n=3; ns: P>0.05; *P < 0.05, **P < 0.01, ****P < 0.0001).

**Figure S3.**
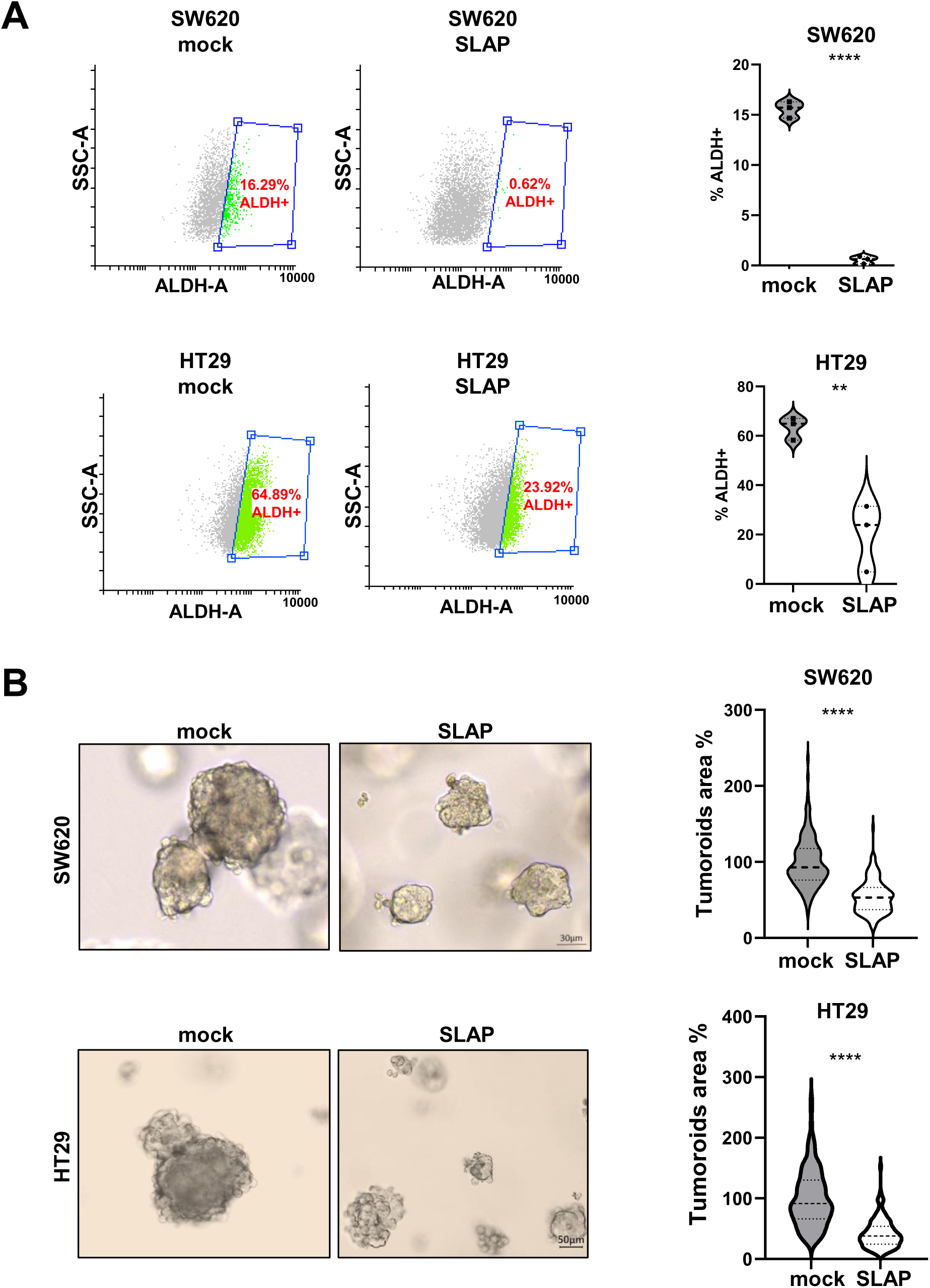
SLAP suppresses human CRC stem cell properties. **(A)** SLAP overexpression reduces ALDH activity in SLAP-low HT29 and SW620 cells (representative FACS plots and quantification; mean ± SEM; n=3; **p<0.01, ****p<0.0001; unpaired t-test).**(B)** SLAP overexpression decreases tumoroid formation in HT29 and SW620 cells (representative images and quantification 25-50 organoids/condition; mean ± SEM; n=3; ****p<0.0001; Mann–Whitney).

**Figure S4.**
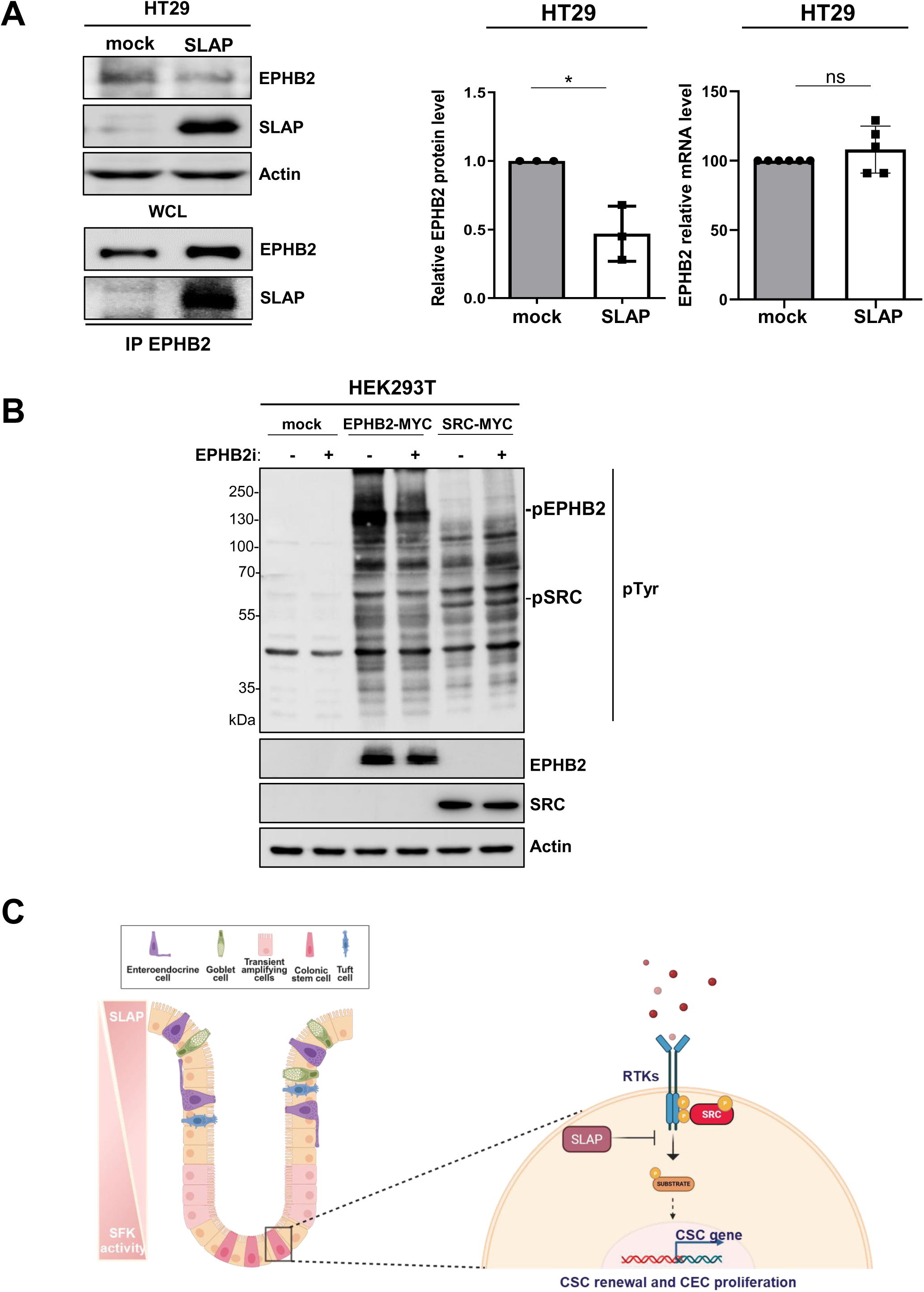
SLAP interacts with EPHB2 and reduces its levels in HT29 cells. **(A)** EPHB2 expression and association with SLAP in CRC cells (transcript and protein; mean ± SEM; n=3-5; ns>0.05; *p<0.05; t-test). **(B)** EPHB2i (400 nM) reduces EPHB2 activity (i.e. pTyr level) in HEK293T cells. **(C)** Model of SLAP regulation of SFK signaling in CSCs to maintain colonic epithelial homeostasis by limiting RTK activity.

**Figure S5.**
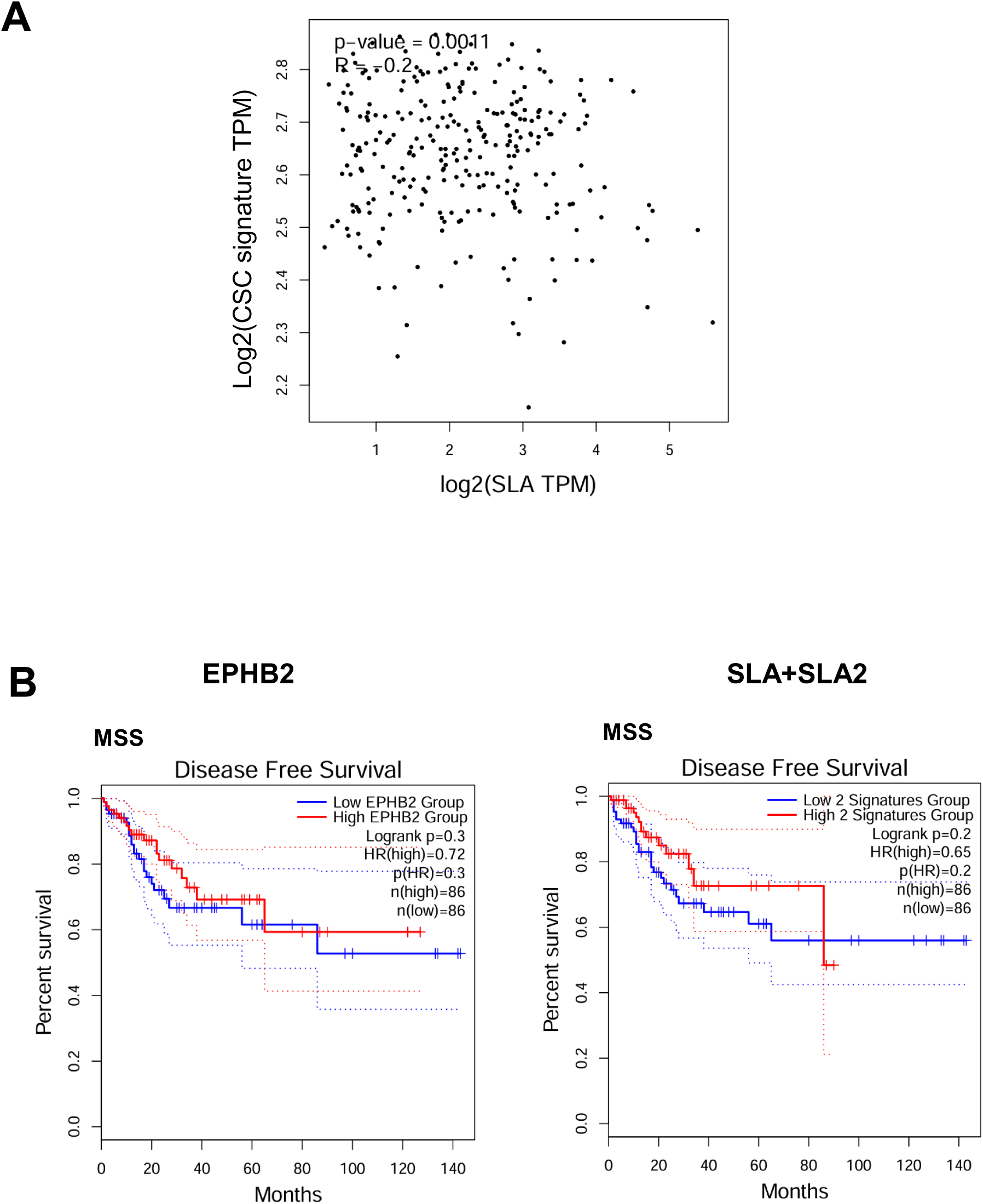
Contribution of SLAP expression to the regulation of EPHB2-dependent CSC signaling in CRC patients. **(A)** Negative correlation between SLAP expression and curated CSC/progenitor gene signature (15 genes including LGR5, ASCL2, OLFM4, BMI1, PROM1, CD44, MKI67, EPCAM, SOX9, SMOC2, TNFRSF19, AXIN2, CYP26A1, ITGA6, PCNA) in TCGA COAD samples (R = −0.2, p = 0.0011). **(B)** EPHB2, or SLA and the SLAP-related gene SLA2 expression do not correlate with disease-free survival in TCGA MSS COAD samples.

